# RsmV a small non-coding regulatory RNA in *Pseudomonas aeruginosa* that sequesters RsmA and RsmF from target mRNAs

**DOI:** 10.1101/315341

**Authors:** Kayley H. Janssen, Manisha R. Diaz, Cindy J. Gode, Matthew C. Wolfgang, Timothy L. Yahr

**Affiliations:** Department of Microbiology, University of Iowa; Department of Microbiology and Immunology, University of North Carolina at Chapel Hill, Chapel Hill, NC; Cystic Fibrosis/Pulmonary Research and Treatment Center, University of North Carolina at Chapel Hill, Chapel Hill, NC

**Keywords:** Pseudomonas aeruginosa, RsmA, RsmF, RsmV, RsmW, RsmY, RsmZ

## Abstract

The Gram-negative opportunistic pathogen *Pseudomonas aeruginosa* has distinct genetic programs that favor either acute or chronic virulence gene expression. Acute virulence is associated with twitching and swimming motility, expression of a type III secretion system (T3SS), and the absence of alginate, Psl, or Pel polysaccharide production. Traits associated with chronic infection include growth as a biofilm, reduced motility, and expression of a type VI secretion system (T6SS). The Rsm post-transcriptional regulatory system plays an important role in the inverse control of phenotypes associated with acute and chronic virulence. RsmA and RsmF are RNA-binding proteins that interact with target mRNAs to control gene expression at the post-transcriptional level. Previous work found that RsmA activity is controlled by at least three small, non-coding regulatory RNAs (RsmW, RsmY, and RsmZ). In this study, we took an in-silico approach to identify additional sRNAs that might function in the sequestration of RsmA and/or RsmF and identified RsmV, a 192 nt transcript with four predicted RsmA/RsmF consensus binding sites. RsmV is capable of sequestering RsmA and RsmF in vivo to activate translation of *tssA1*, a component of the T6SS, and to inhibit T3SS gene expression. Each of the predicted RsmA/RsmF consensus binding sites contribute to RsmV activity. Electrophoretic mobility shifts assays show that RsmF binds RsmV with >10-fold higher affinity than RsmY and RsmZ. Gene expression studies revealed that the temporal expression pattern of RsmV differs from RsmW, RsmY, and RsmZ. These findings suggest that each sRNA may play distinct roles in controlling RsmA and RsmF activity.

**IMPORTANCE:** The role of RsmF in post-transcriptional control of gene expression remains enigmatic. While numerous *rsmA*-dependent phenotypes are more pronounced in an *rsmAF* double mutant, deletion of *rsmF* alone has only modest effects. Understanding mechanisms that control RsmF activity will provide insight into additional roles for RsmF. In the current study we identify RsmV as an sRNA that controls RsmA and RsmF activity, and show that RsmV, RsmW, RsmY, and RsmZ are differentially expressed during growth.

## INTRODUCTION

*Pseudomonas aeruginosa* is a Gram-negative opportunistic pathogen that can cause acute infections in the immunocompromised and chronic infections in individuals with cystic fibrosis (CF) (1, 2). Acute *P. aeruginosa* maladies include skin and soft tissue infections, ventilator associated pneumonia (VAP), and urinary tract infections. *P. aeruginosa* isolated from acute infections are typically motile, non-mucoid, and toxigenic. Acute infections by multi-drug resistant *P. aeruginosa* are difficult to resolve and can progress to sepsis resulting in a high rate of morbidity and mortality (3). Chronic *P. aeruginosa* infections are most common in CF patients and result from a variety of mutations in the CFTR ion channel that result in dehydrated and thickened mucus, and physiochemical changes in the airway surface fluid that result in a clearance defect (4). The persistence of *P. aeruginosa* in the CF airways is associated with adaptive changes including loss of motility, growth as a biofilm, mucoidy, and loss of some acute virulence functions (5, 6). The coordinate transition from an acute to a chronic infection phenotype is regulated by a variety of global regulatory networks including the Rsm system (7).

The Rsm system controls ~10% of the *P. aeruginosa* genome including the type III and type VI secretions, exopolysaccharides important for biofilm formation, and motility (8, 9). The Rsm system includes two small RNA-binding proteins (RsmA and RsmF/RsmN) and at least three small non-coding RNAs (RsmW, RsmY, and RsmZ) that function by sequestering RsmA and RsmF from mRNA targets. RsmA and RsmF are part of the CsrA family and regulate gene expression at the post-transcriptional level. RsmA and RsmF are 31% identical at the amino acid level and both rely on a conserved arginine residue for RNA-binding activity (13, 16). RsmA and RsmF directly interact with mRNA targets to positively or negatively alter translation efficiency and/or mRNA stability (8, 10, 11). The RsmA and RsmF bindings site on target mRNAs commonly overlap the ribosome binding site and consist of a conserved 5’- CAN<u>GGA</u>YG sequence motif (where N is any nucleotide, the underlined GGA is 100% conserved, and Y is either a cytosine or uracil) that presents the GGA sequence in the loop portion of a stem-loop structure (12-15). While RsmA is able to bind mRNA targets with a single CAN<u>GGA</u>YG sequence (12), RsmF differs in that high affinity binding is only observed with mRNAs targets possessing at least two CAN<u>GGA</u>YG consensus binding sites (12). Although RsmA and RsmF share some targets in common, the full extent of overlap between the regulons is unknown (13, 16). The RsmF regulon, however, was recently determined using a pull-down method to identify 503 target RNAs (17).

The RNA-binding activity of RsmA is controlled by the small non-coding RNAs RsmW, RsmY and RsmZ (13, 18). RsmW, RsmY, and RsmZ each have multiple CAN<u>GGA</u>YG binding sites that allow for sequestration of RsmA from target mRNAs (19). It is unclear whether RsmW, RsmY, and RsmZ are the only sRNAs that function in the sequestration of RsmA or whether RsmW, RsmY, or RsmZ are the primary sRNAs that function in sequestration of RsmF. The affinity of RsmF for RsmY and RsmZ is 10-fold lower than RsmA. In this study, we sought to identify additional sRNAs that regulate RsmA and RsmF activity, and identified RsmV, a 192 nt transcript that has four CAN<u>GGA</u>YG sequences presented in stem-loop structures. We demonstrate that RsmV is able to sequester RsmA and RsmF in vivo, that full RsmV activity is dependent upon each of the four CAN<u>GGA</u>YG sequences, and that RsmV demonstrates a temporal expression pattern that is distinct from RsmW, RsmY, and RsmZ. We propose a model wherein each sRNA plays differential and distinct roles in control of the Rsm system.

## RESULTS

### Identification of RsmV as a sequestering RNA for RsmA and RsmF

We took an in-silico approach to identify candidate sRNAs that might control RsmA and/or RsmF activity *in vivo*. A prior SELEX study concluded that optimal RNA-binding activity by RsmF requires RNA targets with at least two GGA sequences presented in the loop portion of stem-loop structures. Transcriptome studies have identified ~500 potential sRNAs in *P. aeruginosa* (28, 29). The secondary structure of each sRNA was predicted using mFold and then examined for the presence of ≥2 GGA sequences presented in stem-loop structures. One sRNA candidate had six GGA sequences (Fig. 1A). Each GGA sequence demonstrated a ≥60% match to the full RsmA/RsmF consensus binding site (CAnGGAyG) (Fig. 1B). Four of the GGA sequences (designated sites 2, 3, 5 and 6) are predicted by mFold to be presented in stem-loop structures (Fig. 1A). The gene encoding the sRNA is located in the intergenic region between *mucE* and *apqZ,* and has been designated *rsmV* (Fig. 1C). A search of the Pseudomonas Genome Database indicates that the *rsmV* sequence is highly conserved in >100 sequenced *P. aeruginosa* genomes and is absent from the genomes of other Pseudomonads.

**Figure 1.**
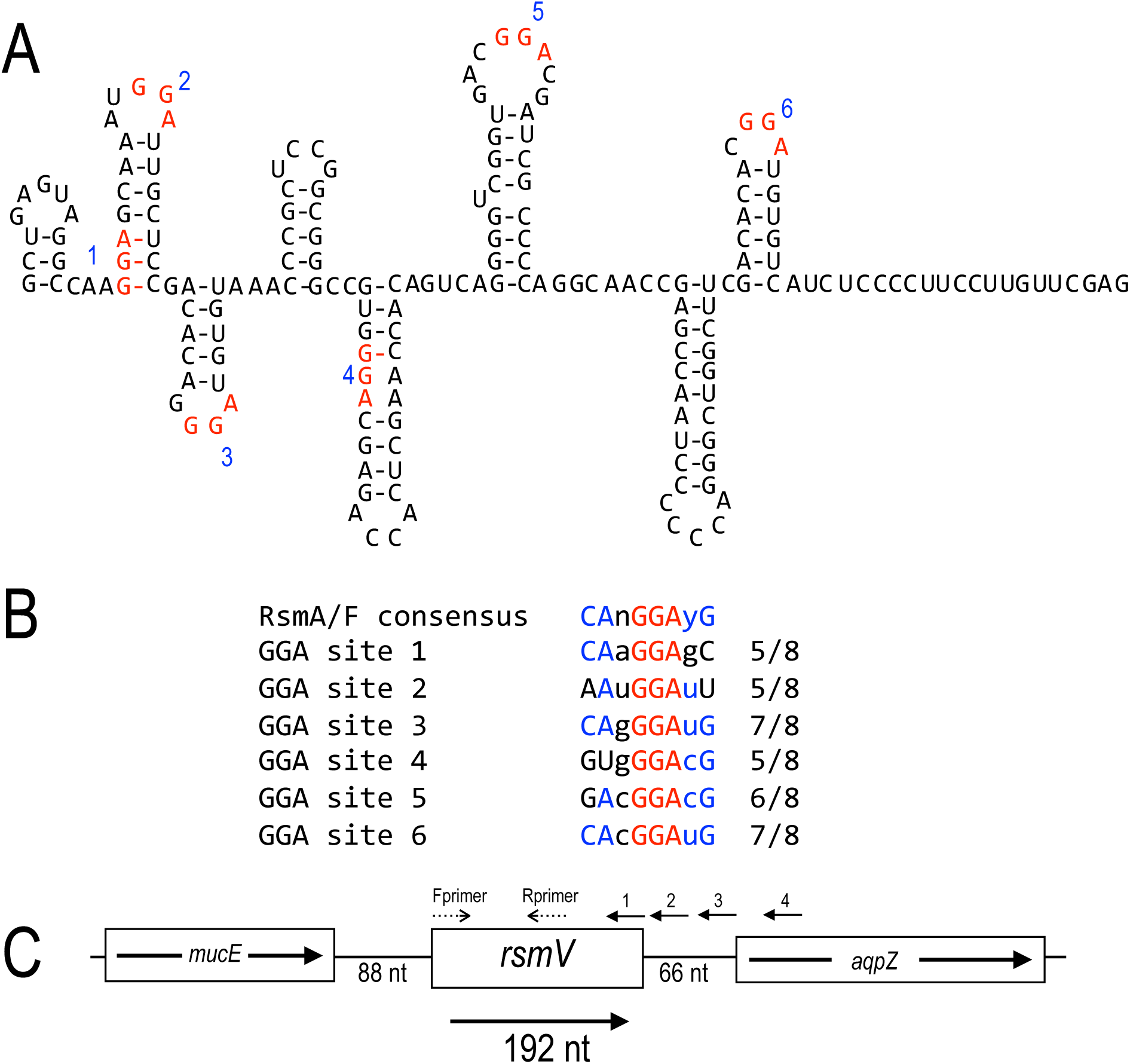
Predicted structure of RsmV and genomic context. (*A*) Predicted mFold structure of RsmV. *P. aeruginosa* RsmV secondary structure determined by mFold modeling. Each of the six GGA sequences are highlighted in red and numbered by order of appearance from the 5’ end of the sequence. (*B*) Alignment of each GGA site to the full RsmA/RsmF consensus binding site. The GGA sites are 100% conserved (red) and other conserved portions of the consensus are highlighted in blue. (*C*) The genome context of *rsmV,* located between *mucE* and *aqpZ*. The positions of the primer used to generate cDNA for the experiment in Fig. 2B are labeled 1- 4. Primers used to generate PCR products in Fig. 2B are labeled Fprimer and Rprimer.

The RNAseq study that identified *rsmV* concluded that the RNA is 192 nt long (29). To verify the *rsmV* transcription start site, cDNA was generated using a primer within the gene (Fig. 2A). The cDNA was then used in a PCR reaction with primers positioned just upstream of and at the predicted start site. Whereas the primer positioned at the start site generated the expected product, the primer located just upstream did not generate a product. This finding is consistent with the *rsmV* transcription start site identified in the Wurtzel RNAseq study (29). There is no identifiable transcriptional terminator downstream of *rsmV*. To verify the 3’ boundary, cDNA was generated from total cellular RNA with primers positioned at the predicted 3’ end of *rsmV* and at several downstream positions as shown in Fig. 1C. The resulting cDNAs were then used as templates in PCR reactions with primers (Fprimer and Rprimer) positioned within the gene (Fig. 1C). The cDNA primer positioned within the *aqpZ* coding region (primer 4) did not yield a product. The cDNA primers positioned upstream of *aqpZ* (primers 1, 2, and 3) all yielded products, and the strongest product was observed with cDNA generated at the predicted 3’ terminus of *rsmV* using primer 2 (Fig. 2B). The weaker PCR products produced from cDNA generated with primers 2 and 3 may represent transcriptional read through.

**Figure 2.**
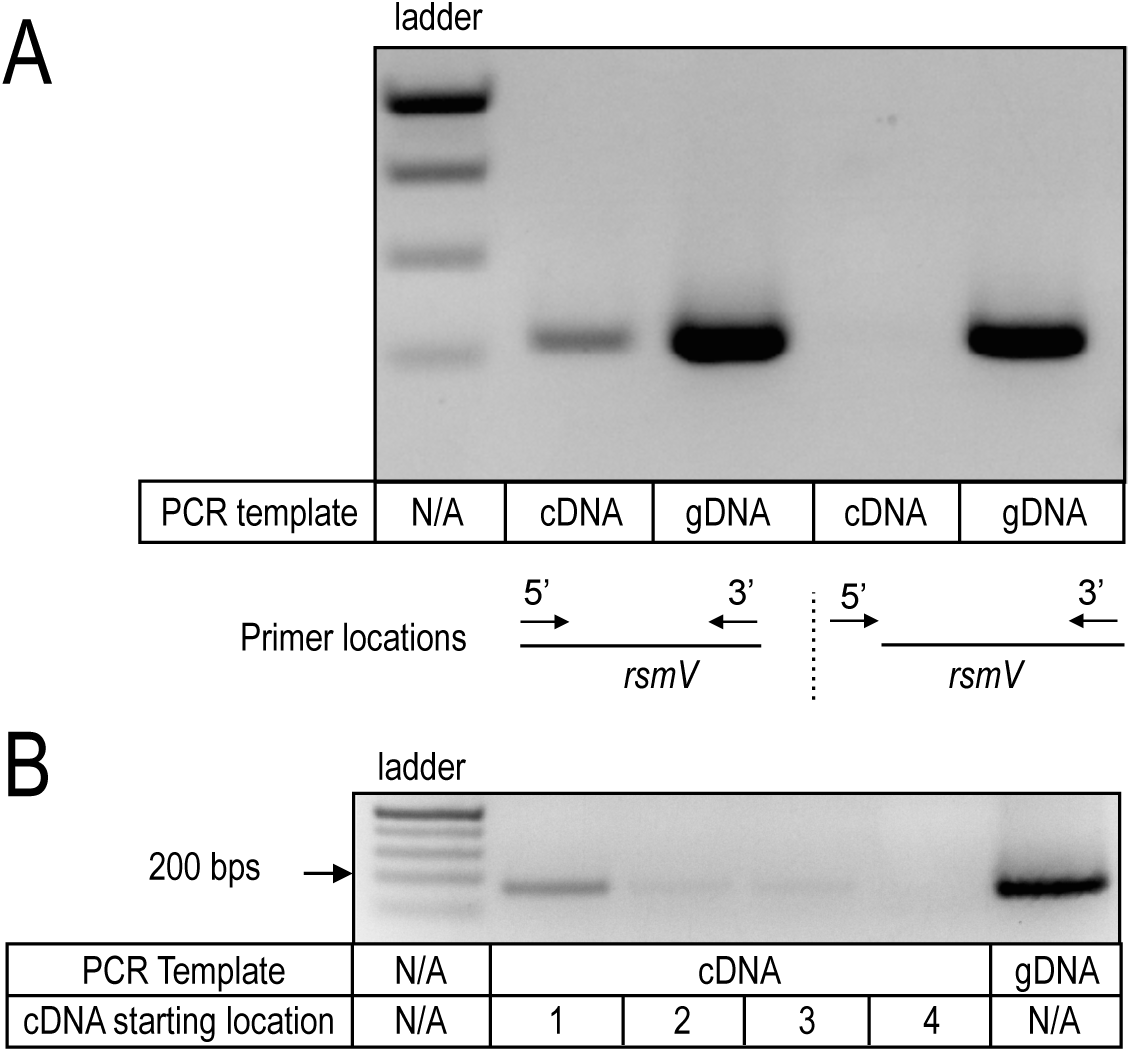
Verification of the *rsmV* 5’ and 3’ boundaries. (*A*) RNA purified from wt cells was used to generate cDNA using the indicated 3’ primer. The cDNA was then used in PCR reactions with the same 3’ primer and 5’ primers positioned just upstream of or at the predicted start of *rsmV* transcription. Genomic DNA (gDNA) served as a positive control. (*B*) Verification of the *rsmV* termination site. cDNA was generated using primers 1-4 as shown in Fig. 1C. The cDNA was then used in PCR reactions with the indicated primer sets in Fig. 1C. Genomic DNA (gDNA) served as a positive control.

### RsmV interacts with and controls RsmA and RsmF activity

The presence of four predicted GGA sequences in stem-loop structures is consistent with RsmV serving as a sequestering sRNA for RsmA and/or RsmF. To test this prediction we measured binding using EMSA experiments. Full length RsmV was synthesized *in vitro*, radiolabeled at the 5’ end, and incubated with purified RsmA_His_ or RsmF_His_ prior to electrophoresis on non-denaturing gels. RsmA_His_ formed high affinity binding products with RsmV (*K*_eq_ 14 nM) and two distinct binding complexes were evident (Fig. 3A). Those products could reflect binding of multiple RsmA_His_ dimers or differential interactions with multiple sites on the RsmV probe. RsmF also bound the RsmV probe with high affinity (*K*_eq_ 2 nM), but only a single binding complex was detected (Fig. 3A).

**Figure 3.**
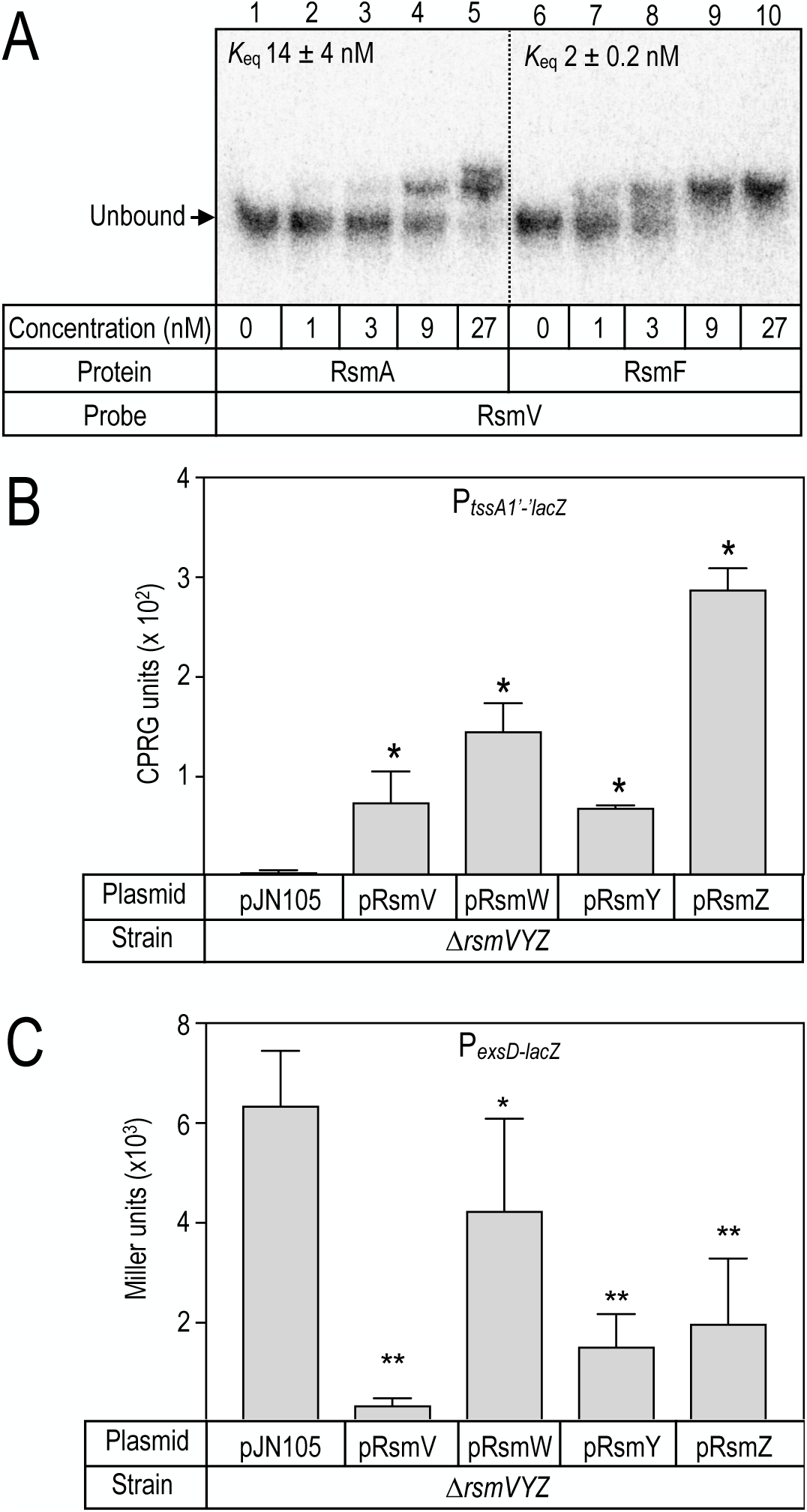
RsmV binding and regulatory activity. (*A*) RsmV was radiolabeled and used in electrophoretic mobility shift assays with purified RsmA (lanes 2-5) and RsmF (lanes 7-10) at the indicated concentrations. The position of the unbound RsmV probe is indicated. (*B-C*) Effect of RsmV on *tssA1’-‘lacZ* translational reporter (*B*) and P*_exsD-lacZ_* transcriptional reporter (C) activities. Strains consisting of a ∆*rsmVYZ* mutant transformed with either a vector control (pJN105) or the indicated sRNA expression plasmids were cultured in the presence of 0.4% arabinose to induce expression of the respective RNAs and assayed for ß-galactosidase activity. The reported values represent the average of at least three experiments with the standard error indicated. * *P*-value <0.05 relative to the vector control.

With evidence that both RsmA and RsmF interact with RsmV we next examined whether RsmV can sequester RsmA/RsmF in vivo. RsmA and RsmF have inverse effects on expression of the type VI (T6SS) and type III (T3SS) secretion systems (13). We used the previously described P*_tssA1’-‘lacZ_* translational reporter as surrogate for regulatory control of T6SS (13) and P*_exsD-lacZ_* transcriptional reporter as marker for the T3SS (30). RsmA/RsmF directly bind the *tssA1* leader region to inhibit translation (12, 13) and positively regulate T3SS gene expression through a mechanism that remains to be defined (13). In a mutant lacking *rsmV*, *rsmY*, and *rsmZ*, RsmA/RsmF availability is high resulting in repression of P*_tssA1’-‘lacZ_* reporter activity and high levels of P*_exsD-lacZ_* reporter activity (Fig. 3B-C). Plasmid expressed RsmV resulted in significant activation of P*_tssA1’-‘lacZ_* reporter activity and inhibition of the P*_exsD-lacZ_* reporter. Both of these findings are consistent with RsmV serving a role in RsmA/RsmF sequestration. When compared to the previously identified sequestering RNAs RsmW, RsmY, and RsmZ (13, 31), RsmV demonstrated activity comparable to RsmY when using the P*_tssA1’-‘lacZ_* reporter (Fig. 3B) and had the strongest inhibitory activity for the P*_exsD-lacZ_* reporter (Fig. 3C).

To determine whether RsmV preferentially sequesters either RsmA or RsmF, the P*_tssA1’- ‘lacZ_* translational reporter was introduced into ∆*rsmAVYZ* and ∆*rsmFVYZ* mutant backgrounds. RsmA is more active than RsmF resulting in stronger repression of P*_tssA1’-‘lacZ_* reporter activity in the ∆*rsmFVYZ* mutant when compared to the ∆*rsmAVYZ* background for strains carrying the vector control (pJN105) (Fig. 4A vs 4B). In the ∆*rsmFVYZ* mutant, where repression of P*_tssA1’-‘lacZ_* activity is attributable to RsmA, RsmV demonstrated relatively weak suppressive activity when compared to RsmW, RsmY, and RsmZ (Fig. 4A). A similar picture emerged when using the ∆*rsmAVYZ* background to examine RsmF sequestration in that RsmV demonstrated the weakest suppressive activity (Fig. 4B). Thus, RsmV is capable of sequestering both RsmA and RsmF and appears to lack strong preference for one vs the other under the conditions tested.

**Figure 4.**
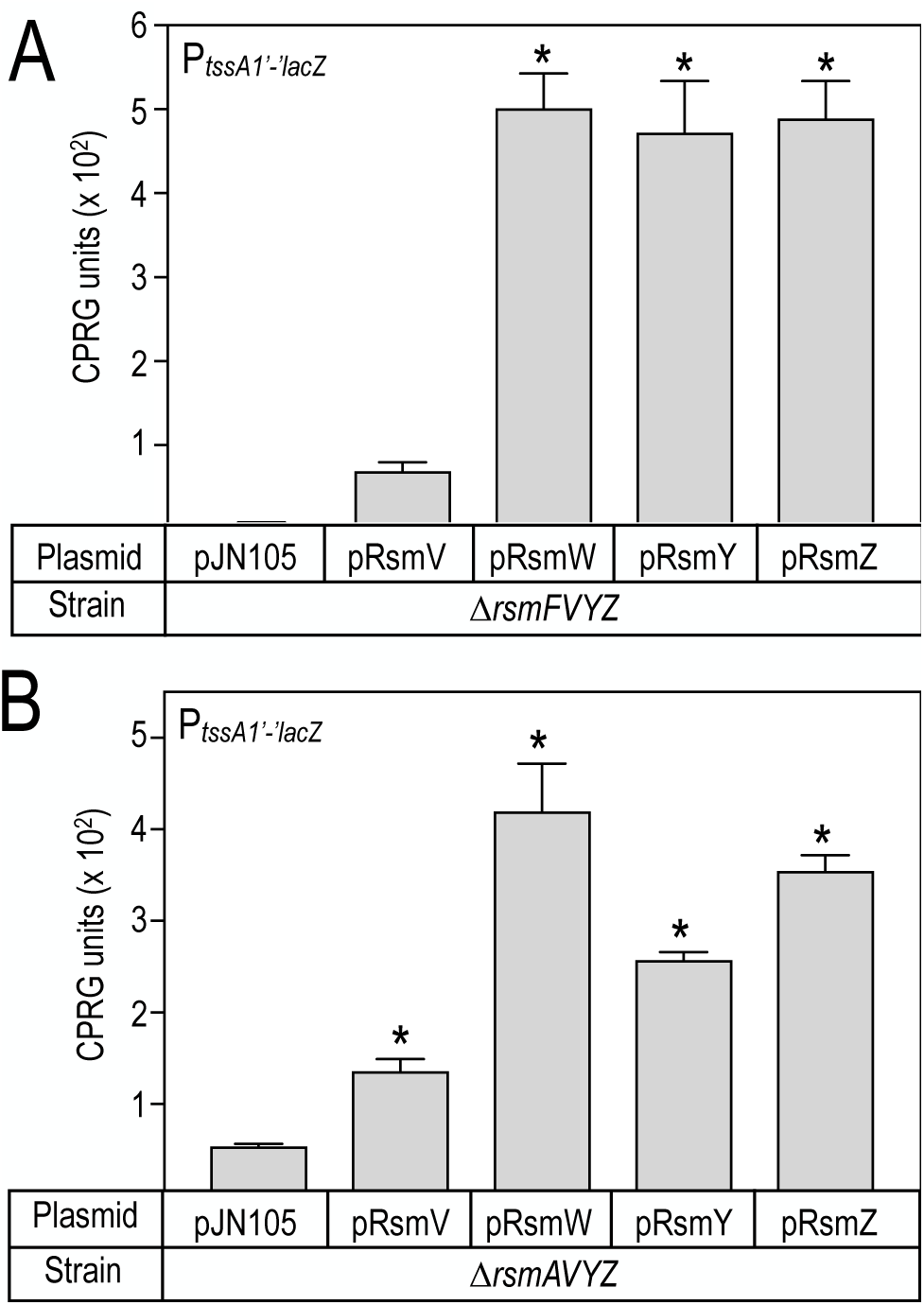
Sequestration of RsmA or RsmF by the RsmV, RsmW, RsmY and RsmZ regulatory RNAs. (*A-B*) Either ∆*rsmAVYZ* (*A*) or ∆*rsmFVYZ* (*B*) quadruple mutants carrying the P_lac-*tssA1*’-‘*lacZ*_ translational reporter was transformed with either a vector control (pJN105) or RsmV, RsmW, RsmY and RsmZ expression plasmids. The resulting strains were cultured in the presence of 0.4% arabinose to induce expression of the respective RNAs and assayed for ß-galactosidase activity. Reported values represent the average of at least three experiments with the standard error indicated. * *P*-value <0.05 relative to the vector control.

### Contribution of GGA sites 2, 3, 5 and 6 to RsmV activity

The RsmV primary sequence contains six GGA sequences, four of which (GGA2, 3, 5, and 6) may be presented in the loop portions of stem-loop structures (Fig. 1A). To determine which GGA sites are important for RsmV regulatory activity each of the GGA sequences in stem-loop structures was changed to CCU. The activity of each mutant RNA was tested using the P*_tssA1’-‘lacZ_* translational and P*_exsD-lacZ_* transcriptional reporters. The GGA4 and GGA6 mutant RNAs demonstrated a significant loss of regulatory activity for the P*_tssA1’-‘lacZ_* translational reporter when compared to wt RsmV (Fig. 5A). In contrast, each of the GGA sites was required for full regulatory control of the P*_exsD-lacZ_* transcriptional reporter (Fig. 5B). The most likely explanation for the differential requirement for the GGA2 and GGA5 sites is that the P*_exsD-lacZ_* reporter is more sensitive to changes in RsmA availability relative to the P*_tssA1’-‘lacZ_* reporter.

**Figure 5.**
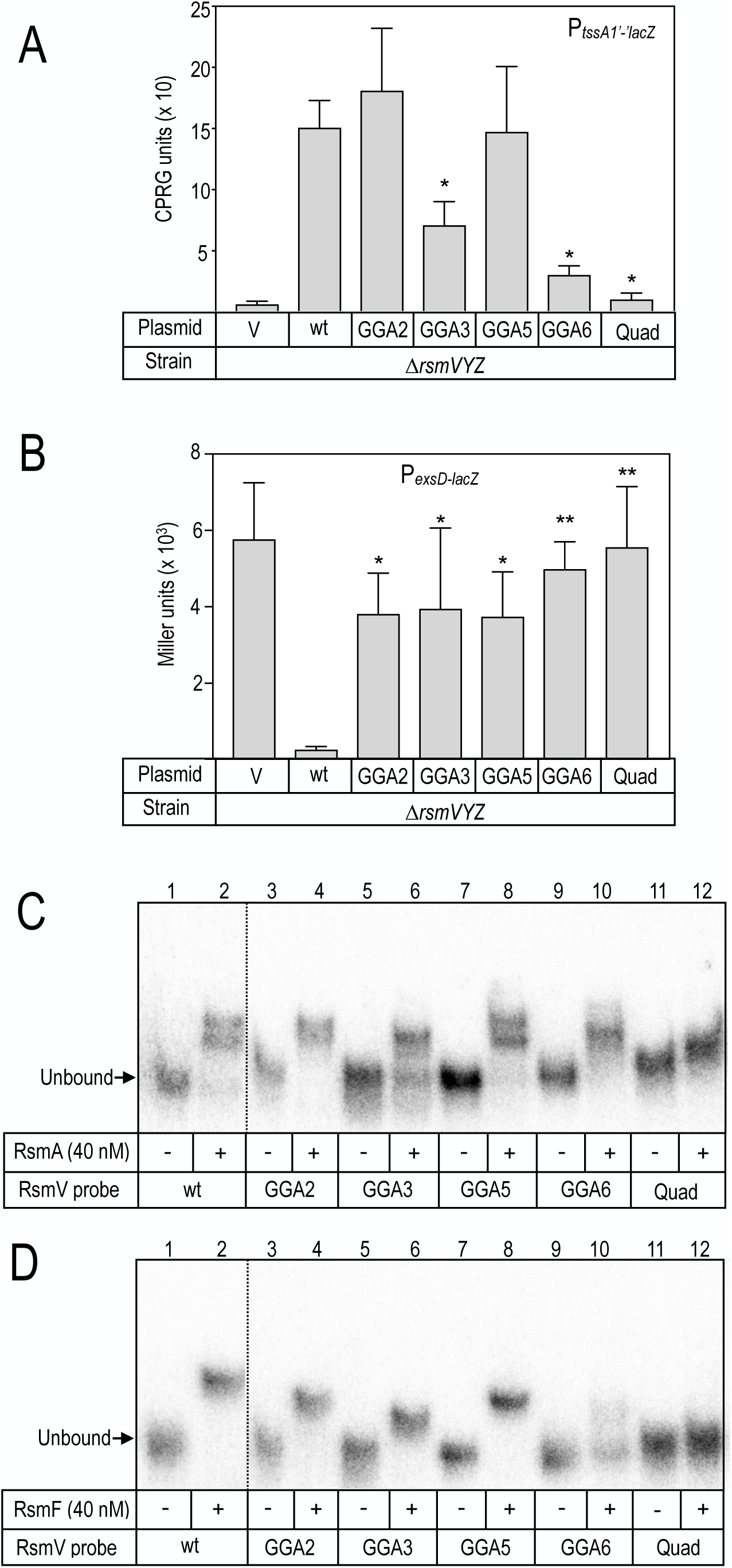
Functional analyses of the RsmV mutants. The PA103 ∆*rsmVYZ* mutant carrying the (*A*) P_lac_ *tssA1*’-‘*lacZ* translational reporter or (*B*) *P_exsD_-lacZ* transcriptional reporter were transformed with either a vector control (pJN105) or the indicated RsmV expression plasmids. The resulting strains were cultured in the presence of 0.4% arabinose to induce expression of the respective RNAs and assayed for ß-galactosidase activity. Reported values represent the average of at least three experiments with the standard error indicated. * *P*-value <0.05 relative to the vector control. (*C-D*) EMSA experiments with wt RsmV and the indicated mutant radiolabeled probes. 40 nM RsmA (*C*) or RsmF (*D*) were incubated with the indicated probes, subjected to non-denaturing gel electrophoresis, and phosphorimaging. The positions of the unbound probes are indicated.

The simplest interpretation of the reporter findings is that the mutant RNAs with altered activity have reduced capacity to sequester RsmA/RsmF. To test this prediction, binding assays were performed with radiolabeled RNA probes. RsmA bound each of the single GGA substitution mutants with affinities similar to or greater than wt RsmV (Fig. 5C, Table 1, Fig. S1). Whereas two distinct products are formed upon RsmA binding to the wt, GGA2, and GGA5 probes, only a single product was observed for the GGA3 and GGA6 probes. This is noteworthy as the mutant GGA3 and GGA6 RNAs also demonstrated a defect in activation of the P*_tssA1’-‘lacZ_* translational reporter (Fig. 5A). RsmF also bound each of the mutant probes with high affinity, with the exception of GGA6, which was significantly reduced (Fig. 5D, Table 1, Fig. S1). Given that RsmA and RsmF are homodimers with two RNA binding sites (one from each monomer), and that each mutant RNA still has three potential GGA interaction sites, high affinity binding to the mutant probes was not unexpected. We thus generated a probe bearing CCU substitutions at all four sites (Quad) and found that RsmA (*K*_eq_ >27 nM) and RsmF (*K*_eq_ >243 nM) were unable to bind (Fig. 5C-D, Table 1, Fig. S1) or exert regulatory control over the P*_tssA1’-‘lacZ_* translational and P*_exsD-lacZ_* transcriptional reporters. We conclude that the primary, if not exclusive, sites for RsmA/RsmF binding are GGA2, 3, 5, and 6.

**Table 1.** RsmA and RsmF affinities for wt and mutant RsmV

| RNA | RsmA | RsmF |
| --- | --- | --- |
| wt RsmV | $14 \pm 4^a$ | $2 \pm 0.2$ |
| GGA2 | $3 \pm 3$ | $1 \pm 0.6$ |
| GGA3 | $1 \pm 0.4$ | $5 \pm 3$ |
| GGA5 | $3 \pm 3$ | $8 \pm 13$ |
| GGA6 | $9 \pm 11$ | $>243$ |
| Quad | $>27$ | $>243$ |

<sup>a</sup> apparent equilibrium binding constant (nM)

### Role of RsmV in vivo

Data presented thus far have relied upon plasmid-expressed RsmV, which may result in RNA levels that exceed the native level expressed by cells under physiologically relevant conditions. To address the effect of RsmV expressed at native levels on the output of the Rsm system we generated an in-frame *rsmV* deletion mutant (∆*rsmV*) and measured P*_exsD-lacZ_* reporter activity. When compared to wt cells, the ∆*rsmV* demonstrated a modest but significant increase in reporter activity (Fig. 6A). This increase in reporter activity is consistent with reduced sequestration of RsmA and/RsmF, both of which have a positive effect on T3SS gene expression. By comparison, P*_exsD-lacZ_* reporter activity is also elevated in an ∆*rsmYZ* double mutant. The higher level of reporter activity in the ∆*rsmYZ* mutant is consistent with the data presented in Fig. 3C showing that RsmY and RsmZ each have stronger effects on activation of P*_exsD-lacZ_* reporter activity relative to RsmV.

**Figure 6.**
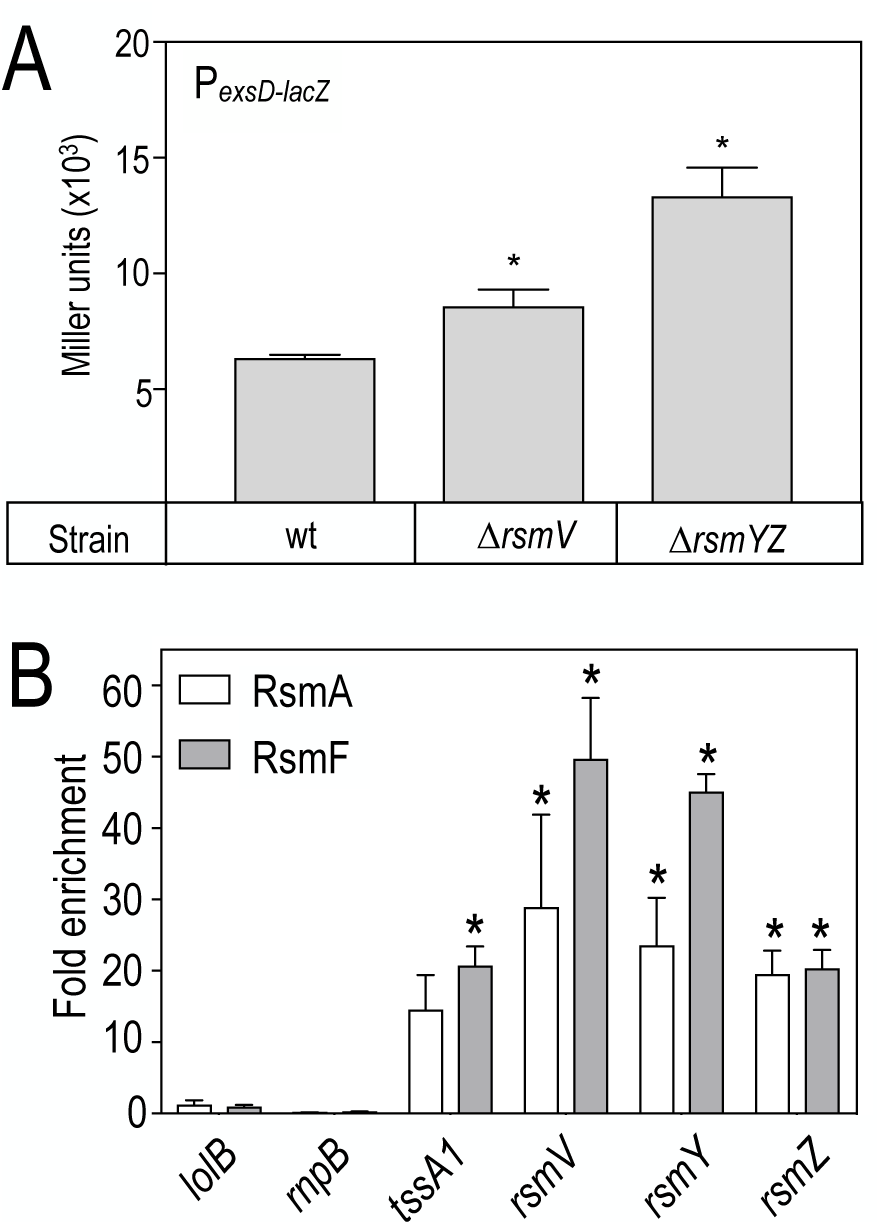
In vivo activity of RsmV. (*A*) Strain PA103 (wt), and the ∆*rsmV* and ∆*rsmYZ* mutants carrying the *P_exsD_-lacZ* transcriptional reporter were cultured under inducing conditions for T3SS gene expression and assayed for ß-galactosidase activity. * *P*-value <0.05 relative to wt. (*B*) A ∆*rsmAF* mutant transformed with either a vector control, or pRsmA_His_ or pRsmF_His_ expression vectors was cultured and subjected to rapid purification of pRsmA_His_ or pRsmF_His_ and bound RNAs. Select RNAs (as indicated) were quantified from the purified RNA pool by qRT-PCR and reported as fold change relative to the vector control. The coding sequences for *lolB* and *rnpB* were included as negative controls and *tssA1* served as a positive control. Reported values represent the average of at least three replicates with the standard error reported. \**P*-value <0.05 when compared to expression of the wild type vector is indicated.

A second approach to test the relevance of RsmV in vivo involved precipitation experiments with histidine-tagged RsmA or RsmF. A ∆*rsmAF* double mutant transformed with either RsmA_His_ or RsmF_His_ expression plasmids was cultured to mid-log phase and then rapidly subjected to precipitation with Ni^2+^-agarose beads and isolation of bound RNA. The presence of specific RNAs was detected from the entire pool of bound RNAs by qRT-PCR. Positive controls were the known RsmA/RsmF targets RsmY, RsmZ, and the *tssA1* leader region (13). Negative controls included two mRNAs (*lolB* and *rnpB*) that are not known targets of RsmA or RsmF, and Ni^2+^-agarose beads alone. Whereas no enrichment of the *lolB* or *rnpB* mRNAs was detected, there was significant enrichment of the *tssA1* mRNA and the RsmV, RsmY, and RsmZ sRNAs by both RsmA and RsmF (Fig. 6B).

### Differential expression of RsmV, RsmW, RsmY, and RsmZ

The in vivo data demonstrate that RsmV, RsmW, RsmY, and RsmZ are each capable of sequestering RsmA and RsmF. We hypothesized that differential expression of the RNAs might allow cells to fine-tune the output of the Rsm system. To test for differential expression, RNA samples were collected from cells cultured to OD_600_ readings of 0.5 (early log phase), 1.0 (mid-log phase), 2.0, 5.0, and 7.0 (late stationary phase). The amount of each RNA detected by qRT-PCR at early log phase was normalized to 1.0 and the reported values for each subsequent time point are relative to those values (Fig. 7). Both RsmY and RsmZ showed a transient increase in expression at mid-log phase, followed by a decrease at OD_600_ readings of 2.0 and 5.0, and then a significant increase in late stationary phase (OD_600_ 7.0) The expression pattern for RsmW was delayed until the OD_600_ reached 2.0 but demonstrated the highest fold changes in expression at OD_600_ 2.0 and 5.0, and then approached the fold changes observed for RsmY and RsmZ at OD_600_ 7.0. By contrast, RsmV demonstrated a slow but steady increase throughout the growth curve but was the least dynamic of the four RNAs. The observed differences in expression patterns are consistent with the hypothesis that the sRNAs may serve distinct roles in RsmA/RsmF sequestration based upon their timing of expression.

**Figure 7.**
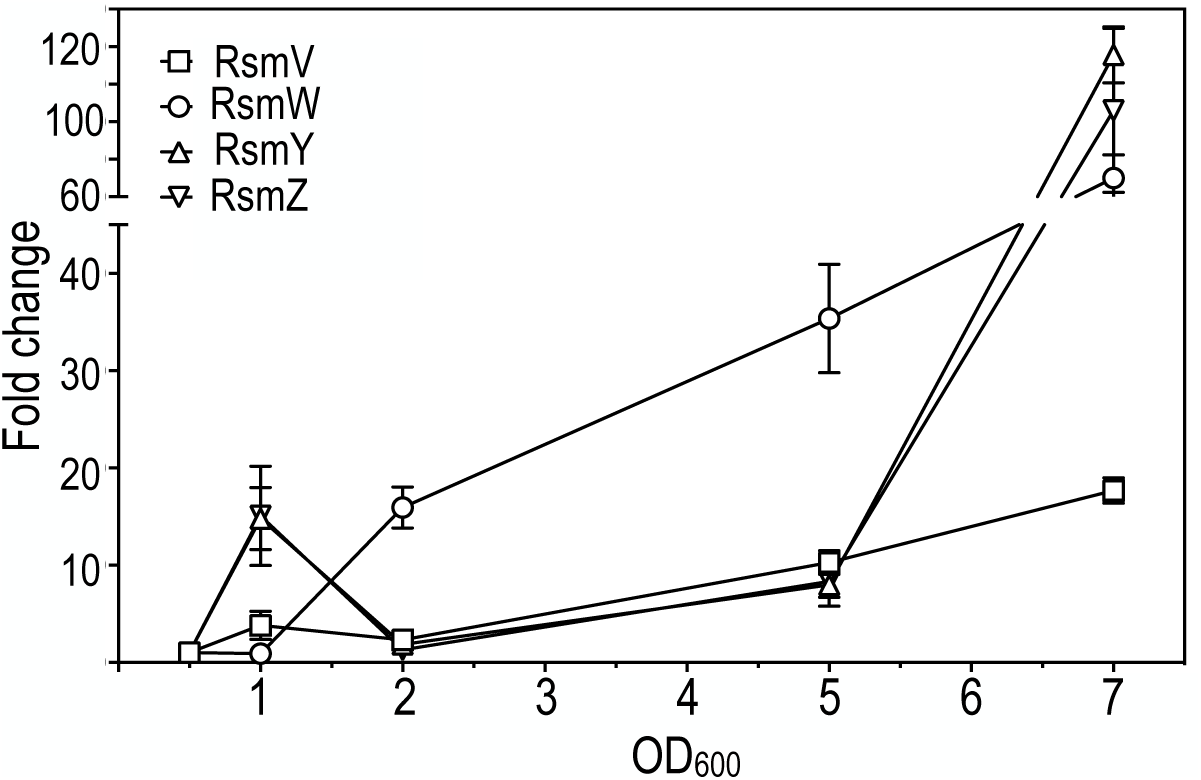
Expression profiles of RsmV, RsmW, RsmY, and RsmZ during a growth curve. RNA was isolated from wt cells harvested at the indicated A_600_ readings and used as template in RT-qPCR experiments with primers specific to *rsmV*, *rsmW*, *rsmY*, and *rsmZ*. The reported values for each RNA are relative to the measurement of the sample collected at A_600_ 0.5. The data represent the average of at least three replicates.

One mechanism to account for the differential expression of RsmV, RsmW, RsmY, and RsmZ is by distinct transcription factors. Transcription of *rsmY* and *rsmZ* is controlled by the GacAS two component system (20). A previous study found that GacAS does not control *rsmW* transcription (31). To determine whether *rsmV* transcription is regulated by GacA, a P*_rsmV-lacZ_* transcriptional reporter was integrated at the ΦCTX phage attachment site of wt cells and a ∆*gacA* mutant. Whereas P*_rsmY-lacZ_* and P*_rsmZ-lacZ_* reporter activity demonstrate strong *gacA*- dependence, P*_rsmV-lacZ_* activity showed no difference between wildtype and the *gacA* mutant (Fig. S2).

## DISCUSSION

The primary RsmA/RsmF sequestering RNAs in *Pseudomonas aeruginosa* are RsmY and RsmZ. In addition to RsmY/RsmZ, RsmW plays a smaller role in the sequestration of RsmA (31) and can also sequester RsmF (Fig. 2). RsmV represents a fourth RsmA/RsmF sequestering RNA in *P. aeruginosa.* RsmV shares sequence and structural characteristics with RsmY and RsmZ including multiple GGA motifs (6), four of which are likely presented in stem-loop structures. RsmY and RsmZ are also the primary sequestering RNAs in *P. fluorescens* (now *P. protegens)* (18). At least one additional sRNA, RsmX, also contributes to Rsm control in *P. protegens* (32). The involvement of multiple sequestering sRNAs in the control of CsrA/RsmA activity is common. CsrB is the primary CsrA-sequestering RNA in *E. coli* and contains 18 GGA motifs (33). Other *E. coli* sRNAs can also sequester CsrA including CsrC and McaS (34, 35). CsrC has a structure similar to CsrB but with fewer GGA motifs (34). McaS is an sRNA that basepairs with some mRNAs involved in curli and flagella synthesis and can also sequester CsrA via two GGA motifs (35). In addition to sRNAs, the 5’ untranslated region of mRNAs can also function in the sequestration of CrsA (36).

The relative activities of RsmV, RsmW, RsmY, and RsmZ were compared by expressing each sRNA from an arabinose-inducible expression vector. Plasmid expressed RsmV activated P*_tssA1’-‘lacZ_* reporter activity and inhibited P*_exsD-lacZ_* reporter activity (Fig. 3B-C). RsmV had activity comparable to RsmY for activation of the P*_tssA1’-‘lacZ_* reporter activity and the strongest effect on activation for P*_exsD-lacZ_* reporter activity. RsmA and RsmF both bind to RsmV with high affinity in vitro (Fig. 3A), and the affinity of RsmF for RsmV is at least 10-fold higher than for RsmY and RsmZ (12, 13). Although the affinity of RsmF for RsmV is higher in vitro, RsmV does not seem to show preferential activity towards RsmF over RsmA in vivo (Fig. 4). The reason for this is unclear but may reflect differences between the in vitro and in vivo binding conditions. A difference between in vitro and in vivo conditions was also evident when the RsmV GGA mutants were examined. Whereas each of single GGA substitution mutants demonstrated altered regulatory control of P*_exsD-lacZ_* and/or P*_tssA1’-‘lacZ_* reporter activity (Fig. 5A-B), the binding affinity of RsmA and RsmF was relatively unaffected by the single GGA substitutions (Table 1). A similar trend was observed in a previous mutagenesis study of RsmY and RsmZ wherein the in vivo activity did not strictly correlate with in vitro binding (37). It was speculated that other RNA binding proteins, such as Hfq, may prevent binding to suboptimal sites in vivo.

RsmV activity is clearly evident when expressed from a plasmid (Fig. 3). A role for RsmV when expressed at native levels from the chromosome was also detected. Deletion of *rsmV* resulted in a modest but significant increase in T3SS reporter activity (Fig. 6A) and co-purification experiments found that RsmV interacts with RsmA and RsmF (Fig. 6B). Both of these findings suggest that RsmV can compete with RsmY and RsmZ for RsmA/RsmF binding in wt cells (Fig. 6B). Unclear is whether conditions exist where *rsmV* transcription is elevated and might result in more pronounced phenotypes. Transcription of *rsmY* and *rsmZ* is directly controlled by the GacA/S two-component system, a highly conserved system in Gamma-proteobacteria (20, 38, 39). GacS is a sensor kinase whose activity is controlled by two orphan kinases, RetS and LadS (40-43). Additional regulators interact with and alter the effect of RetS on GacS (44, 45). SuhB regulates *rsmY* and *rsmZ* transcription indirectly by altering *gacA* levels (24). The phosphotransfer protein, HptB, regulates *rsmY* and *rsmZ* transcription when *P. aeruginosa* is grown on a surface (22, 23). Other regulators contribute to *rsmY* and *rsmZ* transcription through mechanisms that do not alter GacS/GacA activity. MvaT, a H-NS like protein, binds A+T rich regions of DNA and silences *rsmZ* transcription, while BswR, a transcriptional regulator, counteracts negative regulation of *rsmZ* by MvaT (20, 21). Recently, MgtE, a magnesium transporter, was shown to alter *rsmY* and *rsmZ* transcription, yet a mechanism of action is yet to be defined (25).

Neither *rsmV* nor *rsmW* are under positive transcriptional control of the GacAS system (31) (Fig. S2). GacA may repress *rsmW* transcription through an indirect mechanism (30). RsmW expression appears to be highest during stationary phase in minimal media, which may be more biologically representative of a biofilm (31). RsmW is encoded directly downstream of PA4570, a protein of unknown function. RsmW and PA4570 are likely co-transcribed and separated by an RNase cleavage event. Determining the transcriptional regulation of PA4570 may provide insight into *rsmW* transcriptional control. A search for potential promoters upstream of *rsmV* predicted binding sites for the transcriptional activators RhlR, AlgU, and FleQ. mRNA levels for *rsmV,* however, were unaffected in PA14 transposon mutants within each of those genes relative to wild type as measured by qRT-PCR (data not shown). Additional studies will be required to determine how *rsmV* and *rsmW* transcription is controlled, and if RsmV plays a larger role in regulating RsmA and/or RsmF activity under a different set of growth conditions.

RsmX, RsmY, and RsmZ in *P. protogens* are differentially expressed, thus contributing to a mechanism of fine-tuning RsmA and RsmE activity (32). Expression of *P. protogens* RsmX and RsmY occurs in parallel during exponential growth while RsmZ expression is delayed (32). This may allow cells to fine-tune expression of these sRNAs based on the environmental conditions. We propose a similar scenario for expression of RsmV, RsmW, RsmY, and RsmZ in *P. aeruginosa*. The differences in binding affinities for RsmA/RsmF, timing of gene expression, and expression levels of the sRNAs may provide a mechanism of fine-tuning the expression of genes under control of the Rsm system.

## METHODS AND MATERIALS

### Strain and plasmid construction

Routine cloning was performed with *E. coli* DH5*α* cultured in LB-Lennox medium with gentamycin (15 μg/ml) as required. *P. aeruginosa* strain PA103 and the ∆*gacA*, ∆*rsmYZ* mutants were reported previously (Table 1) (47). The in-frame ∆*rsmV* deletion mutant was constructed by allelic exchange. The upstream and downstream flanking regions (~800 bp) of *rsmV* were generated by PCR using primer pairs 118845409- 118845410 and 118845411-118845412. The PCR products were cloned into pEXG2 (48) and the resulting construct was mobilized into wild type PA103 and the ∆*rsmYZ*, ∆*rsmAYZ*, and ∆*rsmFYZ* mutant by conjugation. Merodiploids were resolved by sucrose counter-selection as previously described (49). The RsmV expression plasmid was constructed by positioning the *rsmV* transcription start site immediately downstream of the P_BAD_ promoter start site using the Gibson assembly method (New England Biolabs). Briefly, the P_BAD_ promoter region from pJN105 (primer pair 117830775-117830776) and *rsmV* (primer pair 118845423-118845424) were amplified by PCR and then assembled into the MluI and SacI digested pJN105 (50). pRsmV vectors bearing single GGA to CCT substitutions, or various combinations therefore, were assembled using the Gibson method from gene blocks listed in Table 2 and cloned into the NruI and PvuI sites of pJN105 as outlined in Table 3. The *rsmV* transcriptional reporter (primer pair 150592489-150592490) includes 500 nucleotides upstream of the *rsmV* transcription start site. The *rsmV* reporter was integrated into the CTX phage attachment site in WT and *gacA* strains.

### β-Galactosidase Assays

PA103 strains were grown overnight at 37°C in LB containing 80 μg/ml gentamicin as required. The next day strains were diluted to an absorbance (A_600_) of 0.1 in tryptic soy broth (TSB) for measurement of *tssA1’*-*‘lacZ* reporter activity or TSB supplemented with 100 mM monosodium glutamate, and 1% glycerol for measurement of P*_exsD-lacZ_* reporter activity. Arabinose (0.4%) was also added to induce *rsmV* expression from the P_BAD_ promoter. The cultures were incubated at 37°C and harvested when the A_600_ reached 1.0. β-galactosidase activity was assayed with the substrates ortho-nitrophenyl-galactopyranoside (ONPG) as previously described (51) or chlorophenol red-β-D-galactopyranoside (CPRG). CPRG activity was determined by measuring product formation at 578 nM and using an adaptation of the Miller equation: CPRG units = (A_578_/culture A_600_/time /culture vol [ml]) x 1000. CPRG and Miller units are reported as the average of at least three independent experiments with error bars representing the standard deviation (SD).

### Electrophoretic mobility shift assays

DNA templates encoding wildtype *rsmV* or *rsmV* bearing point mutations within the GGA sequences were PCR amplified and used as templates for in vitro generation of RNA probes. RNA probes were end-labeled with [γ-^32^] ATP as previously described (13). Purified RsmA or RsmF, as described previously (13), were incubated with the RNA probes at the indicated concentrations in 1X binding buffer (10 mM Tris-HCl pH [7.5], 10 mM MgCl_2_, 100 mM KCl), 3.25 ng/μl total yeast tRNA (Life Technologies), 10 mM DTT, 5% (vol/vol) glycerol, 0.1 units RNAse Out (Life Technologies). Reactions were incubated at 37° C for 30 min, and then mixed with 2 μl of gel loading buffer II (Life Technologies) and immediately subjected to electrophoresis on 7.5 % (wt/vol) native polyacrylamide glycine gels (10 mM Tris-HCl pH[7.5], 380 mM glycine, 1 mM EDTA) at 4° C. Imaging was performed using an FLA-7000 phosphorimager (Fujifilm), and analyzed using MultiGuage v3.0 software.

### RNA enrichment experiments

Strain PA14 *ΔrsmAF* carrying either an empty vector control, pRsmA_His6_, or pRsmF_His6_ was grown in TSB supplemented with 20 mM MgCl_2_, 5 mM EGTA, 15 μg/ml gentamicin, and 0.1% arabinose to mid log phase, chilled, and pelleted for immediate lysis. Cells were lysed under native conditions to retain protein structure (Qiagen QIAexpressionist manual native purification buffer recipe) supplemented with 2.5 mM ribonucleoside vanadyl complex (NEB) to inhibit RNAse activity, 1 mg/mL lysozyme, and 0.1% Triton X-100. Lysis was completed by freeze-thaw cycles. Lysates were treated with 10 uL RQ-1 RNase-free DNAse and cleared by centrifugation. An aliquot was removed from the cleared lysate for total RNA isolation and preserved in Trizol, and the remaining lysate was incubated with Ni-NTA agarose at 4°C for 1 hour under non-denaturing binding conditions. Ni-NTA agarose was then loaded into a column and washed 3 times with non-denaturing binding buffer containing 10 mM imidazole. Protein and associated RNAs were eluted in 4 fractions with 250 mM imidazole and 4 fractions with 500 mM imidazole. Protein-containing fractions from the RsmA_His6_, or RsmF_His6_ expressing strains, and an equivalent volume from the vector control strain, were treated with TRIzol (Thermofisher) and RNA was extracted according to the manufacturer’s protocol. RNA was treated with RQ-1 RNase-free DNase and concentrated using a RNA Clean and Concentrator kit (Zymo). First strand cDNA was synthesized using Superscript II (ThermoFisher) according to manufacturer’s protocol with Random Primer 9 (NEB). The copy number of the indicated genes was determined by qPCR using SYBR Green Master Mix (Bio-rad).

### Statistical analyses

One-way ANOVA was performed using Prism 6.0 (GraphPad Software, Inc., La Jolla, CA).

## ACKNOWLEDGEMENTS

This work was supported by the National Institutes of Health, grant number AI097264 to MCW and TLY. KHS was supported by T32GM082729 and 5T32AI007511-19.

